# Enzymes and Substrates Are Balanced at Minimal Combined Mass Concentration *in vivo*

**DOI:** 10.1101/128009

**Authors:** Hugo Dourado, Veronica G. Maurino, Martin J. Lercher

**Author notes:** Corresponding author: Martin Lercher.

## Abstract

A fundamental problem in biology is how cells organize their resource investment. Cellular metabolism, for example, typically involves hundreds of enzymes and metabolites, but it is unclear according to which principles their concentrations are set. Reasoning that natural selection will drive cells towards achieving a given physiological state at minimal cost, we derive a general equation that predicts the concentration of a metabolite from the concentration of the most abundant and costly enzyme consuming it. Simulations of cellular growth as well as experimental data demonstrate that costs are approximately proportional to molecular masses. For effectively irreversible reactions, the cell maximizes its metabolic efficiency by investing equally into substrate and unbound enzyme molecules. Without fitting any free parameters, the resulting model predicts ***in vivo*** substrate concentrations from enzyme concentrations and substrate affinities with high accuracy across data from ***E. coli*** and diverse eukaryotes (*R*^2^=0.79, geometric mean fold-error 1.74). The corresponding organizing principle – the minimization of the summed mass concentrations of solutes – may facilitate reducing the complexity of kinetic models and will contribute to the design of more efficient synthetic cellular systems.

## Introduction

Optimality principles have been used to predict diverse complex cellular properties (1), such as the efficient use of metabolic networks to convert nutrients to biomass (2), the regulation of ribosome content during bacterial growth (3), or the partitioning of membrane occupancy between transporters and the electron transport chain (4). It is likely that in most cells, natural selection has favoured metabolic efficiency, *i.e*., a near optimal cost/benefit relationship for each active biochemical reaction. While the benefit corresponds to the maintenance of a desired reaction rate, the cost lies in the concentrations of the molecules supporting the reaction (5-7).

It is still unclear what factors dominate the costs associated with concentrations of individual proteins or other molecule types. It has recently been argued that the major cost factor of metabolic reactions stems from protein production costs (8-11), including the allocation of cellular resources such as ribosomes and the consumption of ATP and carbon. Other recent publications have stressed the importance of volumetric costs related to the limited solvent capacity of cellular compartments. The summed volume concentration of solutes cannot exceed critical values, beyond which adequate diffusion would break down (5, 6). Based on the limited solvent capacity, it has been proposed that intermediate metabolite concentrations are minimized by natural selection (5, 6, 12). However, the majority of the cellular volume is occupied by proteins, while metabolites account for only a minor fraction; in *E. coli*, proteins outweigh metabolites 16:1. (13) Accordingly, some authors have argued that the solvent capacity limits enzyme rather than metabolite concentrations (14-16), a phenomenon termed macro-molecular crowding. Limiting the total enzyme concentration while maximizing biomass production indeed allows to predict different variants of overflow metabolism, such as the Crabtree and Warburg effects, at least qualitatively (14-16).

While most previous researchers (8-11, 14-16) considered only the cost of enzymes, reaction rates are jointly determined by enzymes and their substrates. Thus, we propose that the action of natural selection on intracellular concentrations can only be appreciated fully when we consider costs incurred through both types of molecules simultaneously; this argument holds both for production costs (reflecting nutrient consumption) and volumetric costs (reflecting consumption of the limited solvent capacity).

## Results

To examine the consequences of this hypothesis on the balance between substrate and enzyme concentrations for individual reactions, we first consider an enzyme that converts a single substrate irreversibly into a product following Michaelis-Menten kinetics, where the substrate is not consumed by any other reaction. The reaction rate *v* is then proportional to the concentrations of unbound enzyme (*E*_free_) and substrate (*M*_free_), parameterized by the enzyme’s turnover number *k_cat_* and Michaelis constant *K_m_* (inversely related to the enzyme’s substrate affinity): 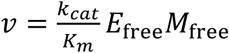. The concentration of unbound enzyme molecules is approximated as a function of the total concentrations of enzyme (*E*) and substrate (*M*) as 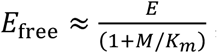; with *M*_free_ ≈ M, rearrangement results in the standard Michaelis-Menten equation.

A reaction rate *v* per unit volume “demanded” by the current cellular state can be achieved in multiple ways: for each non-zero substrate concentration *M*, there is exactly one enzyme concentration *E* (and a corresponding *E*_free_) that results in the desired rate (Figure 1). The costs associated with the enzymatic reaction will depend approximately linearly on *E* and *M* at least for small concentration changes (11); they can thus be expressed through the specific costs of enzyme and substrate molecules, *c_E_* and *c*_*M*_, respectively. Under natural selection for metabolic efficiency, the cell is expected to choose the combination of concentrations that minimizes the summed costs of enzyme and substrate. Because the reaction rate depends equally on the concentrations of unbound enzyme and metabolite, the optimally efficient metabolic state invests equally into these two types of molecules:

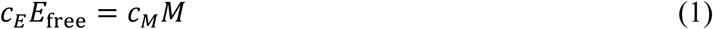

(for a formal derivation and generalizations, see SI Text). Converting to total enzyme concentration *E* and defining the cost ratio of enzyme and substrate molecules *a:*= *c_E_/c*_M_, we can rewrite this as

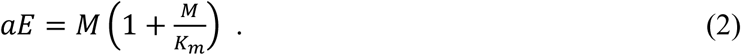

Strikingly, this optimal relationship between enzyme and substrate concentration depends only on the Michaelis constant *K_m_* and on the cost ratio *a*, but is independent of reaction rate and turnover number.

**Figure 1.**
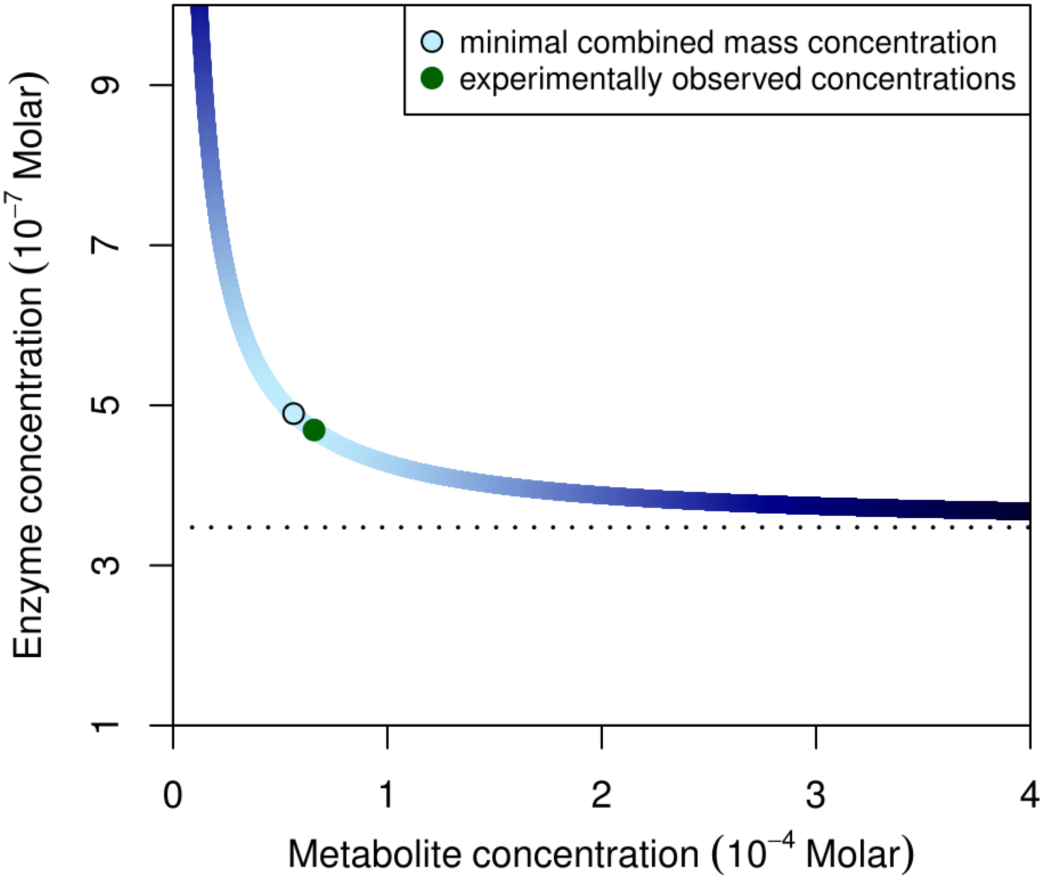
For each Substrate concentration *M*, there is one enzyme concentration *E* that supports a given reaction rate *v*. The curve shown is for the GMP reductase enzyme (GuaC) and its substrate GMP in *E. coli*, color-coded according to the summed mass concentration of GuaC and GMP. The combination resulting in the smallest summed mass concentration is indicated by the circle; the green dot indicates the *in vivo* combination observed for *E. coli* growing on glycerol (23, 24).

Reality is of course more complex than the irreversible single-substrate reaction discussed so far. Cellular metabolism forms a highly interconnected network: many reactions involve more than one substrate, and many metabolites are consumed by more than one enzyme. When considering metabolic costs, we must therefore account for all metabolite and enzyme concentrations simultaneously. Using convenience kinetics to approximate general reaction kinetics (17), it is easy to show that we can consider each metabolite separately as long as the relevant reactions are effectively irreversible (SI Text). The resulting equation for a single metabolite relates its concentration to the concentrations of all enzymes consuming it, mediated by the corresponding Michaelis constants and the relative costs. If one of the enzymes dominates this mathematical relationship, the terms corresponding to the other enzymes can be neglected; typically, the “dominant” enzyme has the largest *aE*, *i.e*., the highest total cost among all enzymes consuming the metabolite. This approximation results in an equation identical to Eq. (2) that considers only the dominant enzyme (SI Text). Note that the dominant enzyme in our terminology may not be the enzyme responsible for the highest turnover of the metabolite, but is the one with the highest total cost in the current metabolic state.

Generalizations to multifunctional enzymes, stoichiometries other than 1:1, reversible reactions, and Hill kinetics are derived and summarized in the SI Text. All considered cases are well approximated by Eq. (2) or its generalization to other stoichiometries.

To compare Eq. (2) to experimental data, we first need to determine the cost ratio *a* of enzymes and metabolites. Indirectly, such costs can be observed as growth rate reductions in experiments that force bacteria to overexpress unneeded proteins or metabolites (7, 10, 18, 19). However, these observations cannot be directly transformed into cost estimates: forced overexpression may cause major cellular reorganizations (20), and the reported effects are unlikely to represent metabolically efficient states (21).

To nevertheless explore different approximations to the enzyme–metabolite cost ratio, we utilize a minimalistic *in silico* cell model (15). Our model is comprised of a small set of transport and enzymatic reactions that follow Michaelis-Menten kinetics and convert two nutrients (termed C and N) to precursors for the production of proteins, including a ribosome, and of lipids (Fig. S1a; for detailed methods see SI Text). The simulations search for a combination of protein and metabolite concentrations that maximizes balanced growth, where all cellular components are reproduced in proportion to their concentrations. This model fully accounts for molecular production costs as well as a solvent capacity limit on the summed volume concentration of all intracellular solutes. Model parameters are the external nutrient concentrations, kinetic constants of the biochemical reactions, protein and metabolite masses, and the cytosolic solvent capacity. To simulate a limiting nutrient, we considered 1000 random parameterizations with low external concentrations of N while allowing unlimited uptake of C.

We first tried to approximate cost ratios through the relative molecular yield of enzyme and metabolite production from the limiting nutrient, *a_N_*, reflecting direct production costs. This resulted in a Pearson correlation of R^2^=0.78 between the simulated metabolite concentrations and those predicted from enzyme concentrations via Eq. (2) (Fig. 2B; geometric mean fold-error *GMFE*=1.83, considering only molecules that require N for their production). For comparison, we repeated this analysis using molecular weights as a proxy for the relative costs of enzymes and metabolites, *a_m_*. Molecular weight appears a reasonable first approximation for production costs of different molecule species, even if this measure does not account for differences in atomic composition, cofactor utilization, and length of required production pathways. Moreover, molecular volumes correlate strongly with molecular weights (22), and thus *a_m_* directly reflects relative volumetric costs. Employing the mass ratio *a_m_* in Eq. (2) resulted in significantly improved predictions (Fig. 2A; R^2^=0.85, *GMFE*=1.68; empirical P-value for the superiority of predictions from *a_m_* compared to *a_N_*: P<10^-15^). Strikingly, the superiority of molecular mass as a proxy for total cost becomes most pronounced when we consider only those metabolites made up exclusively of the limiting nutrient N (*a_m_: GMFE*=2.52; *a_N_*: *GMFE*=4.06). Simulations where C and N are equally growth limiting lead to very similar results (Fig. S1).

**Figure 2.**
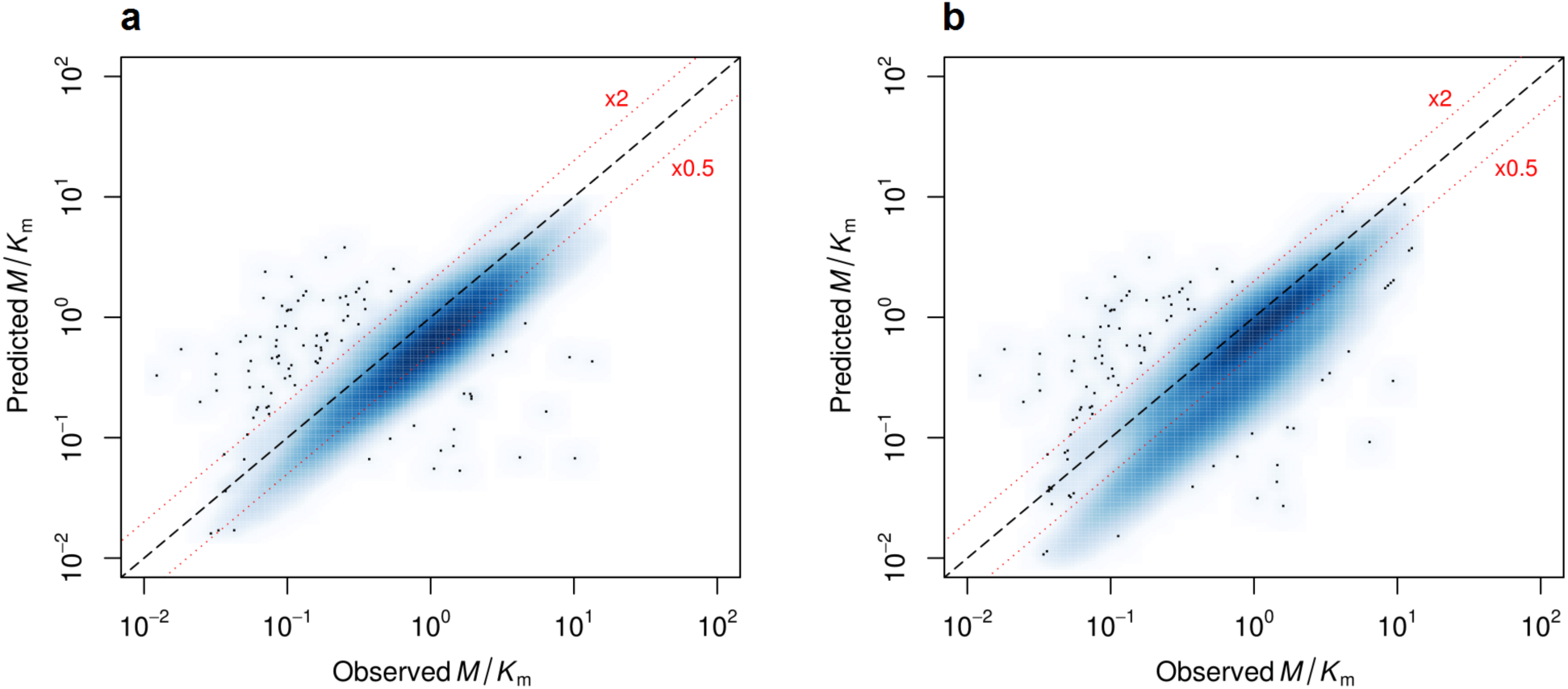
Eq. (2) predicts metabolite concentrations from enzyme concentrations observed in the minimal *in silico* cell accurately when relative enzyme/metabolite costs are approximated through the mass ratio *a_m_* (*R*^2^=0.85, *GMFE*=1.68) (a), but less accurately when approximated through molecular yields for production from the limiting nutrient (*R*^2^=0.78, *GMFE*=1.83) (b). Each smoothed scatter plot summarizes results across all intracellular reactions for 1000 random model parameterizations; colour intensity is proportional to data density, points shown individually are outliers. The black dashed lines indicate the expected identity; the upper and lower red dotted lines indicate deviations of ×2 and ×0.5, respectively.

We conclude that the behaviour of intracellular concentrations in the *in silico* cell is well described by Eq. (2) when approximating relative costs through mass ratios. Can the same model also predict the relationship between enzyme and metabolite concentrations *in vivo?* Experimental data for absolute intracellular concentrations are only available for a limited number of enzyme–metabolite pairs in *Escherichia coli* (23, 24) and in the yeast *Saccharomyces cerevisiae* (25, 26), and for isolated reactions in red blood cells (27) and in the green alga *Chlamydomonas reinhardtii* (28). Application of Eq. (2) requires knowledge of *K_m_* for the dominant enzyme, further reducing the sample size.

All available data accurately conforms to Eq. (2) (Fig. 3a,c). Dominant enzyme concentrations together with *Km* values and molecular mass ratios are sufficient to predict *in vivo* metabolite concentrations with a Pearson correlation of *R*^2^=0.79 (P<10^-16^) and a geometric mean fold-error *GMFE*=*1.74* across the combined *E. coli* and eukaryotic data (Fig. 3b,d). We conclude (i) that the costs of intracellular concentrations can be approximated through molecular masses not only in the *in silico* cell but also *in vivo*, and (ii) that biological cells optimized for metabolic efficiency balance their enzyme and metabolite concentrations accordingly. The metabolite concentrations (26) in yeast were measured for a slightly different strain and on a different growth medium compared to the protein concentration measurements (25); this discrepancy may contribute to the larger deviation between predictions and measurements in the yeast data compared to other cell types (Fig. 3). Both the *E. coli* and the *S. cerevisiae* data includes tRNA-charging reactions (Fig. S2), emphasizing the applicability of the proposed relationships also to non-metabolic enzymatic reactions.

*E. coli* protein expression patterns may not be geared towards maximal metabolic efficiency in many conditions (29). When considering only the *E. coli* data for the two carbon sources most likely to be optimized for metabolic efficiency – those with the highest growth rates, glucose and fructose (23) – we obtain *R*^2^=0.90 (*P*=10^-8^) and *GMFE*=1.51, compared to *R*^2^=0.79 (*P*=10^-16^) and *GMFE*=1.67 when considering *E. coli* data across all growth conditions. Indeed, for most reactions, enzyme concentrations show little variation across conditions, while metabolite concentrations are much more variable (Fig. S2). This might indicate that for each of these reactions, enzyme levels are optimally regulated for growth on glucose and/or fructose regardless of the condition. This pattern would be consistent with an evolutionary strategy that shortens the lag-phase at the transition from an unpreferred to a preferred carbon source, minimizing associated changes in protein expression at the cost of non-optimal expression in unpreferred conditions (30).

**Figure 3.**
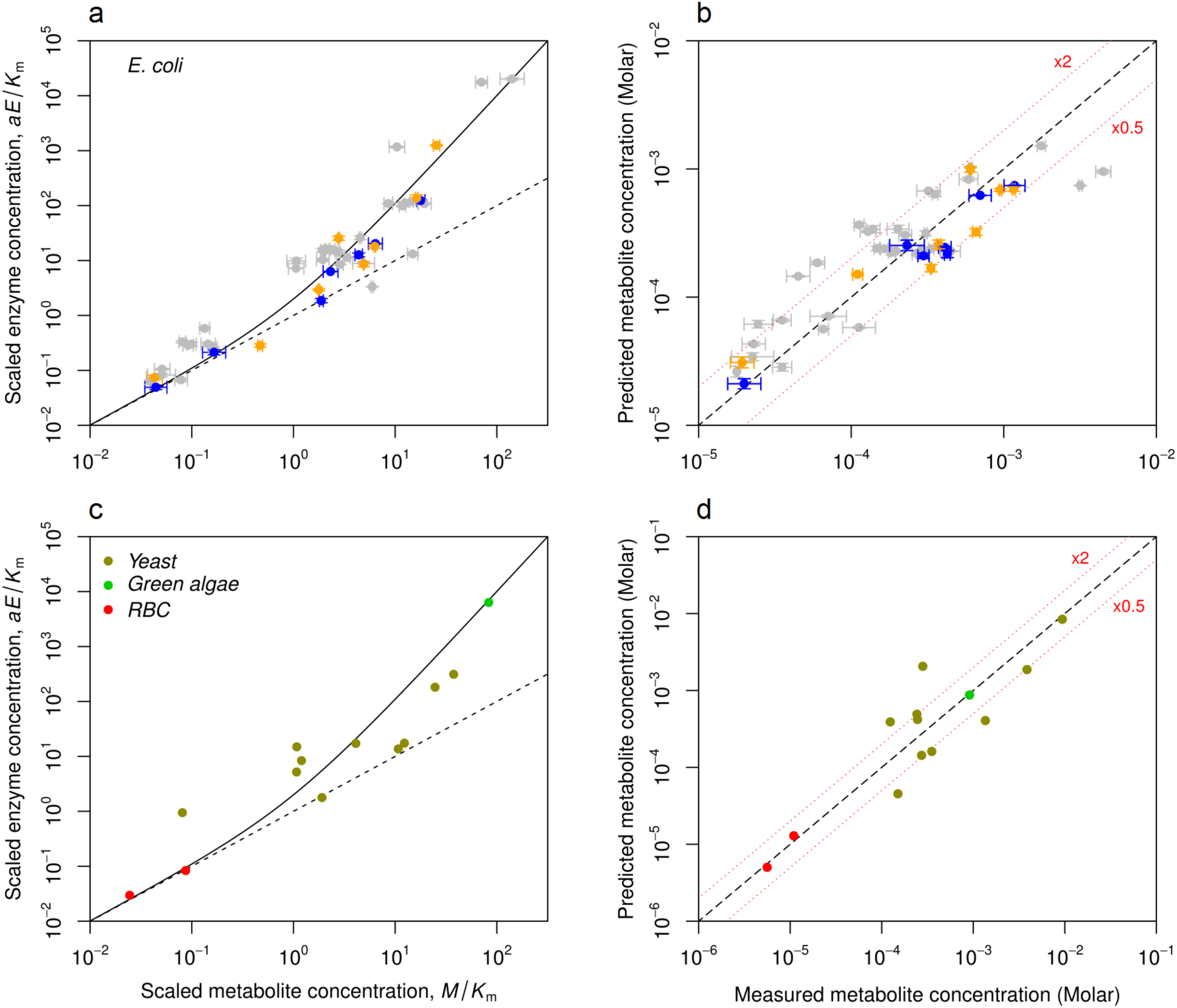
(a) Eq. (2) holds accurately across available data from *E. coli* (orange: growth on glucose; blue: growth on fructose; grey: growth on other carbon sources). Concentrations of metabolites (24) and enzymes (23) are scaled as *x*=*M/K_m_* and *y*=*a_m_E/K_m_* with molecular mass ratio *a_m_*, such that all predictions follow *y* = *x* (1+*x*) (solid line). The diagonal (dotted line) indicates equal cellular mass concentrations of metabolite and enzyme (*a_m_ E* = *M*). Error bars are reported measurement errors. (b) Predicted *vs*. observed *E. coli* metabolite concentrations (Pearson’s correlation between log-transformed values *R*^2^=0.79, *P*=10^-16^; *GMFE*=1.67; data restricted to growth on glucose and fructose: *R*^2^=0.90, *P*=6×10^-8^, *GMFE*=1.51). The dashed line indicates the expected identity. The upper and lower red dotted lines indicate deviations of ×2 and ×0.5, respectively. For the x-axis, error bars are reported measurement errors; for the *y*-axis, error bars reflect the expected standard deviation based on reported measurement errors for enzyme concentrations. See Fig. S2 for the *E. coli* raw data and individual growth conditions. (c) Eq. (2) holds accurately across available data from diverse eukaryotes (yellow: yeast (*Saccharomyces cerevisiae*); red: red blood cell; green: green alga (*Chlamydomonas reinhardtii*)). Concentrations are scaled as in panel (a). (d) Predicted *vs*. observed eukaryotic metabolite concentrations (*R*^2^=0.80, *P*=4x10^-5^; GMFE=2.06).

Red blood cells have little room for enzymatic reactions, as hemoglobin makes up 97% of their cellular dry weight (31). Consequently, metabolite concentrations are much below *K_m_* in this cell type (27), and reaction rates are reduced to 2–8% of their values at saturation. As predicted by the asymptotic behaviour of Eq. (2), the total cellular mass of an enzyme and its substrate becomes very similar in this case (*a_m_E≈M;* Fig. 3c): when most enzymes are in their unbound state, optimal investment is distributed equally across enzyme and substrate molecules. At the other end of the spectrum, Rubisco molecules in the green alga *C. reinhardtii* are highly saturated (*M*=83×K_m_) and outweigh their substrate ribulose 1,5 bisphosphate (RuBP) 75:1, (28) a relationship still accurately predicted by Eq. (2). While all other enzyme–substrate pairs considered here correspond to cytosolic reactions, Rubisco is located in algal chloroplasts, emphasizing the applicability of our results to cellular compartments other than the cytosol.

Several of the reactions included in Fig. 3 are reversible (Table S1), and it is not obvious that Eq. (2) can be applied in these cases. For one of the reactions, Fumarase A (fumA) consuming fumarate (FUM) in *E. coli*, we also have concentration measurements for the product (L-malate) as well as the full kinetic constants of the reversible Michaelis-Menten equation (32). In all assayed conditions, the ratio of product/substrate concentrations is much smaller than the equilibrium constant for this reaction, K_eq_=11.0 (Table S1). As long as the product is not strongly saturating the enzyme, which is the case for the majority of conditions 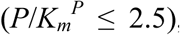, the predictions derived from Eq. (2) do not differ substantially from those derived with its equivalent for reversible reaction kinetics, Eq. (S50). During growth on succinate, however, the product is strongly saturating 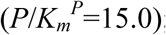; here, employing the reversible prediction reduces the mean fold-error from 4.71 to 1.22 (Fig. S3). The overall excellent match between observed metabolite concentrations and those predicted via Eq. (2) (Fig. 3) suggests that the FUM-fumA case is representative for reversible reactions in general: across most growth conditions, generally reversible reactions may be rendered effectively irreversible by metabolite concentrations that provide an adequate thermodynamic driving force. This notion agrees with previous considerations of the effect of thermodynamic driving forces on expected metabolite concentrations (11, 12), and with a detailed analysis of the relationship between metabolite concentrations and measurements of forward to backward flux ratios (26).

Our hypothesis of natural selection on minimal cellular reaction costs makes two specific predictions when considering one metabolite across different growth conditions. First, if the dominant enzyme remains the same across conditions, then we expect the corresponding points to follow the prediction line from Eq. (2), with different positions corresponding to differences in the flux through the reaction across conditions. This can be seen for galactose-1-phosphate uridylyltransferase (GalT): this enzyme is expressed at very low levels except in growth on galactose, where its substrate shows a correlated increase (Fig. S2e). Second, if an enzyme is dominant for a given substrate only in some of the assayed conditions, we expect to see strong deviations from Eq. (2) in those conditions where other enzymes that consume the same substrate become more costly (have higher *a_m_E*). This can also be observed in the *E. coli* dataset. 1-phosphofructokinase (FruK) is dominant for beta-D-fructofuranose 1-phosphate (F1P) during growth on fructose and predicts the observed F1P concentration within a factor of 1.71. However, FruK contributes only 2.0% and 2.6% to the total enzyme mass consuming this substrate on acetate and glucose, respectively. According to the multi-enzyme relationship (Eq. (S25) of the SI Text), we thus expect the intracellular substrate concentration to be higher than what would be “predicted” from FruK concentrations alone. This is indeed what we find: the observed concentrations are 4.0-fold and 12.7-fold higher, respectively, than “predicted” from Eq. (2) (Table S1).

## Discussion

Estimating reaction costs through the total mass of the molecules involved can only provide a rough approximation to the real cellular expenditure. It is likely that for some metabolite-enzyme pairs, specific biophysical or biochemical constraints lead to further deviations from our predictions. First, natural selection may favour lower concentrations for toxic metabolites than those predicted here. Second, the concentration of metabolites that enter the cell through passive diffusion cannot exceed extracellular concentrations. Third, as examined in the Fumarase A example above, the optimal concentrations of metabolites according to Eq. (2) may provide an insufficient thermodynamic driving force for some reversible reactions, and the generalization given in Eq. (S50) of the SI Text will be more appropriate. Finally, we emphasize that our predictions will only hold for cells whose transcriptional regulation of enzymes and transporters has been optimized by natural selection for metabolic efficiency in the specific condition examined.

Because the same reaction rates can be achieved with different combinations of enzyme and metabolite concentrations (Fig. 1), current metabolic models that account for reaction kinetics are either highly underdetermined or must be constrained by experimental data (33). The organizing principle exemplified by Eq. (2) – a minimal summed mass concentration of solutes – has the potential to massively reduce the complexity of such models, as it provides an objective function capable of distinguishing between alternative kinetic solutions (11). For enzymatic subsystems composed of effectively irreversible reactions, Eq. (2) and its generalizations provide one-to-one correspondences between reaction rates and dominant enzyme as well as metabolite concentrations; the substrate concentration of a non-dominant enzyme can be treated as a constant set by the substrate’s dominant reaction, resulting again in a one-to-one correspondence between reaction rate and enzyme concentration. Such improved kinetic modelling, together with a general appreciation of the cellular organizing principle of minimal summed mass concentrations, will allow the design of more efficient cellular systems and will thereby facilitate further progress in synthetic biology (34).

It is commonly assumed that *in vivo* enzyme and metabolite concentrations are not governed by a general optimization principle, but are determined by the biochemical properties of each metabolite and of the enzymes by which it is consumed (12, 35, 36); reaction rates are often assumed to be governed by enzyme concentrations alone (2, 9, 14, 16). In contrast, theoretical considerations of optimal enzyme properties (37) and large-scale modelling studies (11, 33) as well as metabolite (38) and flux (26) measurements and perturbation experiments (21) indicate that reaction rates are jointly determined by enzyme and metabolite concentrations. Our results explain and quantify this relationship: metabolite and dominant enzyme pools are tightly balanced according to a simple organizing principle, likely reflecting natural selection for the parsimonious use of cellular resources.

## Materials and Methods

### E. coli data

We obtained *Escherichia coli* strain BW25113 enzyme concentrations (23) and metabolite concentrations (24) from two recent publications. Metabolite concentrations in μmol/gCDW (24) were converted to cytosolic molar concentrations based on the same conversion factor between cytosol volume and cell dry weight (2.3 ml/gCDW) used by the original authors (24). Enzyme concentrations in protein mass/cell (23) were converted to cytosolic molar concentrations based on: (i) protein molecular weight (23); (ii) cell dry weight (CDW) estimated based on the relationship PDW/CDW=-0.27869μ+0.64034, derived from Table S1 in Ref. (39), with growth rate μ (in units of 1/h) and total protein dry weight (PDW) measured in each condition (23); (iii) the same conversion factor between cytosol volume and cell dry weight (2.3 ml/gCDW) used for the metabolite concentrations (24). Metabolite molecular weights were obtained from EcoCyc (40).

We collected the Michaelis constants (*K_m_*) of wild-type enzymes from EcoCyc (40), BRENDA (41), and UniProt (42). All experimental values are from *E. coli*, with the exception of four metabolite-enzyme pairs where only data from other organisms are available: D-ribulose 5-phosphate–ribose-5-phosphate isomerase A (Ru5P–rpiA), 1,3-bisphospho-D-glycerate–phosphoglycerate kinase (13DGP–pgk), ADP–phosphoglycerate kinase (ADP-pgk), and glycerone phosphate–fructose bisphosphate aldolase (DHAP-fbaA); we did not consider *K_m_* values of the extremophile *Sulfolobus solfataricus*, as these were obtained from measurements at 70°C. If more than one *K_m_* was listed across the databases (Table S1), we first checked if these values were mostly within the same order of magnitude (*i.e*., if the geometric standard deviation was ≤10); in this case, we used the geometric mean of all available values. Otherwise, we considered the available data for *K_m_* to be too unreliable to be included. The data for pairs of metabolites and dominant enzymes is listed in Table S1.

### S. cerevisiae data

We obtained metabolite concentration data from Ref. (26), in which *Saccharomyces cerevisiae* derived from prototrophic strains S288C and W303 were grown at 30 °C in 2% w/v glucose medium containing Yeast Nitrogen Base (YNB) without amino acids. Enzyme concentration data is from Ref. (25), in which *S. cerevisiae* strain BY4741 was grown at 30 °C in 2% w/v glucose supplemented with YPD medium (with amino acids). The Enzyme concentrations in molecules/cell were converted to cytosolic molar assuming cytosol volume of 21 fl, based on: (i) average cell volume of 42 fl when growing on YPD medium Ref. (43); (ii) cytosol volume is about half of the cell volume Ref. (44). Metabolite molecular weights were obtained from BioCyc. Michaelis constants *K_m_* were collected from Ref. (26), as they are the geometric mean of all available values in BRENDA for *S. cerevisiae;* we confirmed that in each case, the geometric standard deviation was <10, i.e., all reported values were of the same order of magnitude. In almost all cases were no *K_m_* value from *S. cerevisiae* was available, there were also no measurements from other organisms. The *K_m_* for fumarate-URA1 (dihydroorotate dehydrogenase) is missing from BRENDA and from Ref. (26), and was obtained from BioCyc instead. The data for pairs of metabolites and dominant enzymes is listed in Table S1.

The genomes of the yeast strains S288C/W303 and BY4741 are highly similar (45), so that their intracellular concentrations of enzymes and metabolites are likely to be comparable if assayed under the same conditions. However, the two growth media employed by Park *et al*. (26) and Kulak *et al*. (25) differ substantially: while YNB (26) is a defined medium that contains no amino acids, YPD (25) contains peptone and yeast extract, making it rich in amino acids. The different growth conditions are likely to induce different intracellular concentrations of enzymes and metabolites, especially of those molecule types involved in amino acid synthesis. Accordingly, we expect to see more deviations between predicted and measured concentrations in yeast than in the other cell types examined; however, no better matching absolute enzyme and metabolite concentration data is available.

### Red blood cell data

We obtained molecular weights, number of binding sites, and binding site concentrations of enzymes, as well as metabolite concentrations for red blood cells from Ref. (27). Albe *et al*. (27) considered fructose 6-phosphate to be the substrate of pgi; we changed this to glucose 6-phosphate in agreement with the direction of glycolysis. Enzyme concentrations were determined dividing the binding site concentration by the number of binding sites of each enzyme. *K_m_* values were obtained from BRENDA. The values are listed in Table S1.

### Green alga data

We obtained the molar concentration of Rubisco binding sites and its substrate ribulose-1,5-biphosphate (RuBP) in the green alga *Chlamydomonas reinhardtii* from Ref. (28). We calculated the Rubisco molar concentration dividing the reported binding site concentration by the number of binding sites according to BRENDA. We considered the concentrations during steady-state in the wild type cell under constant low light intensity (28), using the geometric mean for RuBP concentration (which was measured twice). The molecular weight of *C. reinhardtii* Rubisco was obtained from BRENDA, and the *K_m_* for RuBP from Ref. (46). The values are listed in Table S1.

### Identification of dominant enzymes in E. coli

For an automated identification of dominant enzymes, we used the sybil and sybilSBML (47) packages in R (48), with the EcoCyc metabolic model for *E. coli* exported as an SBML file using Pathway Tools 19.5 (49). For each metabolite measured in Ref. (24), we first identified all reactions using it as a substrate according to the metabolic model. The gene-reaction associations given in the EcoCyc model through b-numbers were used to map the reactions to the proteins measured in Ref. (23), identified by P-numbers. The concentration of enzymes was determined from the protein concentrations and the enzyme protein stoichiometries obtained from Ref. (50) (assuming stoichiometries of 1 for enzymes not listed in (50)).

For each substrate assayed in Ref. (24), we determined a dominance score (hereafter referred to simply as “dominance”) for each enzyme consuming it that was assayed in Ref. (23). The dominance of an enzyme was defined as the fraction it contributes to the total mass concentration of all assayed enzymes using the substrate. An enzyme was considered “dominant” for the substrate if its dominance was >0.5, *i.e*., its molecules constituted more than half of the total protein mass consuming the substrate. We only attempted to assess dominance if more than half of the enzymes consuming a given substrate were assayed in Ref. (23). We excluded membrane-bound and periplasmic enzymes based on Gene Ontology annotations (51) (GO categories 0016020 (membrane), 0005886 (plasma membrane), 0005887 (integral component of plasma membrane), 0042597 (periplasmic space)), as in these cases the estimated enzyme concentrations will not correspond to actual cytosolic concentrations. If the reaction catalyzed by the dominant enzyme was reversible according to the EcoCyc model, this substrate–enzyme pair was only considered further if the flux through the reaction was measured in the corresponding direction in Ref. (24). Cyclic AMP (cAMP) was not included in the analysis, as the major role of cAMP is not metabolic. cAMP regulates transcription through varying concentrations of cAMP-CPR; accordingly, the only enzyme using it as a substrate (cAMP phosphodiesterase) is unlikely to have a major impact on cAMP concentrations.

### Identification of dominant enzymes in S. cerevisiae

We determined dominant enzymes in *S. cerevisiae* using the same automatic procedure as for *E. coli*, using the Yeast v. 7.6 model Ref. (52) (yeast.sourceforge.net) in SBML format. For each metabolite measured in Ref. (26), we first identified all reactions using it as a substrate according to the metabolic model. The concentration of enzymes was determined from the protein concentrations in Ref. (25), assuming stoichiometries of 1 for enzyme complexes.

For each substrate assayed in Ref. (26), an enzyme was considered “dominant” if its dominance was >0.5. We only attempted to assess dominance if more than half of the enzymes consuming a given substrate were assayed in Ref. (25). We excluded membrane-bound and periplasmic enzymes based on Gene Ontology annotations (51) (GO categories 0016020 (membrane), 0005886 (plasma membrane), 0005887 (integral component of plasma membrane), 0042597 (periplasmic space)), as in these cases the estimated enzyme concentrations will not correspond to actual cytosolic concentrations. For the same reason, we only considered the enzymes dominant if they are located in the “cytoplasm” compartment in the metabolic model and assigned as a component of “cytoplasm” in the Yeast Genome Database Ref. (45). If the reaction catalyzed by the dominant enzyme was reversible according to the BioCyc (53) *S. cerevisiae* model (the Yeast v. 7.6 model doesn’t contain information about reversibility), this substrate–enzyme pair was only considered further if the flux through the reaction was measured in the corresponding direction in Ref. (26).

### Identification of dominant enzymes in red blood cells and green algae

We used the HumanCyc (54) database to identify the enzymes involved in consuming each metabolite measured in Ref. (27). High-throughput enzyme MS/MS measurements and molecular weights for human red blood cells were obtained from Ref. (55) (Table S1). As in the *E. coli* and *S. cerevisiae* analysis, we attempted to determine dominance only for those substrates for which more than half of the consuming enzymes were assayed. Enzymes were considered dominant if their dominance score was >0.5. According to the BioCyc database, RubisCO is the only enzyme consuming D-ribulose-1,5-bisphosphate (RuBP) in *Chlamydomonas reinhardtii*, so it is the dominant enzyme by default.

### Empirical P-value

To test if the predictions for the *in silico* cell model are statistically significantly better when using the molecular mass ratio *a_m_* than when using molecular yield for the limiting nutrient, N, we estimated an empirical *P*-value as follows. As a null model, we assumed that both sets of predictions come from the same distribution (*i.e*., both predictions are equally good). We randomly re-assigned the two predictions using *a_m_* and *a_N_* for each observed data point to two groups and calculated the difference in geometric mean fold-error (*GMFE*) between the groups; this was repeated n=10’000 times. In all of the 10’000 repetitions, the *GMFE* difference was smaller than the *GMFE* difference observed in the *in silico* cell simulations (0.15). The randomized *GMFE* differences were normally distributed. Accordingly, we estimated the *P*-value based on the mean value (2.32×10^-6^) and the standard deviation (2.19×10^-4^) of the randomized *GMFE* differences. The z-score of the *GMFE* difference observed in the *in silico* cell simulations is 68, and thus P<10^-15^.

### Generation of Figure 1

The reaction rate *v* of the GMP reductase reaction (GMP–guaC) in *E. coli* growing on glycerol (green dot in Fig. 1) was calculated through the corresponding Michaelis-Menten equation (Eq. (S1)), assuming saturation of the enzyme with the other substrates, NADPH and H+, the experimentally determined concentrations of enzyme (4.69×10^-7^ Molar) (23) and metabolite (6.58×10^-5^ Molar) (24), and the kinetic parameters *k_cat_*=0.28s^-1^, *K_m_*=2.30×10^-6^Molar obtained from the EcoCyc and BRENDA databases (the *K_m_* value is the geometric mean over the available values, SI Table 1). The optimal enzyme concentration (circle in Fig. 1) was calculated using Eq. (2) with the GMP/guaC molecular weight ratio a_m_=413.76. The curve corresponds to Eq. (S3), with the colour code representing the summed mass concentration of GMP and guaC (Eq. (S4) with molecular weights *m_E_*=149454.2 Da, *m_M_*=361.21 Da).

## Acknowledgments

Deniz Sezer shared important insights into the interpretation of Eq. (2). We thank Peer Bork, Oliver Ebenhöh, David Heckmann, Markus Kollmann, Tabea Mettler-Altmann, Balazs Papp, Daniel Rickert, and Itai Yanai for helpful discussions. This work was supported by the German Research Foundation (DFG grants IRTG 1515 supporting HD; FOR 1186 to VGM; EXC 1028 to VGM and MJL; CRC 680 to MJL).

